# Genome reannotation and effector candidate identification in *Meloidogyne chitwoodi* through gland-specific transcriptome analysis

**DOI:** 10.1101/2025.03.27.645669

**Authors:** Marcella Teixeira, Itsuhiro Ko, Sapinder Bali, Paulo Vieira, Thomas R. Maier, Thomas J. Baum, Cynthia Gleason

**Author notes:** Simplot Plant Sciences, J. R. Simplot company, Boise, ID United States of America. Equal contribution (Marcella Teixeira, Itsuhiro Ko, Sapinder Bali).

## Abstract

The root-knot nematode (RKN) *Meloidogyne chitwoodi* is a threat for potato production in the western United States, negatively impacting potato yield and product value*. Meloidogyne chitwoodi* produce proteins, called effectors, in their esophageal glands that are secreted during parasitism and play integral roles in plant-nematode interactions. Because the esophageal glands are the main effector secretory organ, we performed juvenile gland isolation and gland transcriptome analysis. The data allowed us to improve the *M. chitwoodi* genome annotation. Additionally, the gland-specific transcriptome data gave us an enrichment of gland-localized genes, which was validated by *in situ* hybridization. The gland transcriptome analysis led to the identification of 111 effector candidates. One of the effectors, Mc15g003960, which was highly expressed in the pre-parasitic J2 gland tissue, was further characterized. Expression of Mc15g003960 in Arabidopsis resulted in increased galling by *M. chitwoodi*. However, the ectopic expression of Mc15g003960 in planta did not suppress defense-related callose deposition, suggesting that this effector might be involved in processes other than interfering with plant basal defense responses. Our data shows that using the gland transcriptome, good quality genome annotation and stringent criteria, we can increase the efficiency of effector identification, which can be used to develop more sustainable management tools.

**Authors summary:** The root-knot nematode *Meloidogyne chitwoodi* is a major problem for potato farmers in the western U.S., reducing crop yield and quality. These nematodes produce special proteins, called effectors, in their esophageal glands, which help them infect plants. Since these glands are the main source of effectors, we isolated them from juvenile nematodes and analyzed their gene expression. This helped us improve the nematode’s genome map and identify genes specific to the glands. From this study, we found 111 potential effector genes. One of them, *Mc15g003960*, was highly active before the nematode started feeding. When we introduced this gene into *Arabidopsis* plants, the nematodes caused more damage, but it didn’t seem to weaken the plant’s basic defense system. This suggests *Mc15g003960* is not suppressing plant defenses and has a different role in helping the nematode with successful infection. Overall, our approach helped us identify key effectors more efficiently, which could lead to better ways to manage nematode infestations in the future.

## Introduction

Root-knot nematodes (RKN, *Meloidogyne* spp.) cause substantial damage to crop plants, amounting to millions of dollars in agricultural losses each year [1]. *Meloidogyne chitwoodi* is a RKN that is endemic to the potato-producing regions of Oregon, Idaho, and Washington, where over half of the USA potatoes are grown [2, 3]. This species is problematic for potato growers because the nematodes can infect both the roots and tubers. The tuber infections result in potatoes with rough, bumpy appearance and brown spots under the skin, and there is little industry tolerance for such potato blemishes. Moreover, *M. chitwoodi* is considered a quarantine pest in certain export markets for potatoes grown in the USA, and their presence in a shipment bound for such a market could mean that the entire shipment is rejected [4]. There is currently no genetic resistance available in commercially available potato cultivars [5]. To develop new strategies of *M. chitwoodi* management, we need information about how the nematodes are successful at parasitism, which calls for investigation into nematode parasitism genes, also known as effector genes.

Nematode effector genes encode proteins that facilitate successful infections, either by suppressing the plant immune responses and/or by altering the plant cell metabolism to form the nematode feeding site [6]. The expression of effector genes may occur at any life stage. Following embryogenesis and initial development in the egg stage, the RKN will hatch as second-stage juvenile (J2). This is the motile infective life stage that must enter the plant root and migrate to the vasculature to establish its feeding site. As a result, the J2 releases effector proteins, which include cell wall-degrading enzymes, such as cellulases and polygalacturonases. Once inside the root, the J2 induces root galling and will choose between 6-10 plant cells to convert into its feeding sites called giant-cells [7]. The plant cells that form into giant-cells undergo synchronous waves of mitotic activity uncoupled from cytokinesis, and they are transcriptionally active, resulting in enlarged, multinucleate giant-cells that the nematodes rely on for nutrition. After initially feeding, the J2 will quickly molt 3 more times into the adult stage. The adult female will feed on the giant-cells and produce eggs in a gelatinous matrix on the surface of the root. Because nematode parasitic success requires root penetration and giant-cell formation, the effectors secreted during early stages of parasitism, in the pre-parasitic nematode J2 and during early feeding site formation (parasitic J2), are critical [8]. The pre-parasitic J2 faces the challenge of penetrating the roots and migrating towards the vascular cylinder without triggering insurmountable plant defense responses. In addition, the effectors must suppress any defenses, such as those inadvertently triggered by the pathogen associated molecular patterns (PAMPs), such as ascaroside #18 [9]. These basal plant defenses, known as PAMP-triggered immunity (PTI), are counteracted by nematode effectors that specifically inhibit PTI responses [8, 10].

The identification of J2 effectors is not only interesting from the viewpoint of understanding nematode parasitism, but these effectors could also be targets for gene silencing as a possible future nematode control strategy. As a result, we have focused our current efforts on the pre-parasitic J2 transcriptomes. Previously, we studied the motile *M. chitwoodi* J2 and sedentary parasitic J2 life-stages and found 2,248 genes up-regulated in expression in parasitic nematodes in potato roots at 8 days post-inoculation (dpi) compared to the motile, pre-parasitic J2 life stage [11]. Of these 2,248 genes, 269 contained a signal peptide (SP) sequence and no transmembrane (TM) domain(s). These two characteristics are key criteria used in effector discovery pipelines because they suggest the genes encode secreted proteins. While bioinformatic predictions based on the presence of a signal peptide sequence and absence of transmembrane domains can help identify potential effectors, additional confirmation is typically needed. This is usually done through in situ hybridization (ISH) assays. The ISH assays are performed for each gene of interest to determine whether the transcript is localized to the secretory organ. Most nematode effectors are found to be made in the esophageal glands and then secreted out of the nematode stylet into the plant [12]. Thus, the presence of an ISH probe signal localized in the glands of the nematode is indirect confirmation of secretion. Unfortunately, such assays are laborious, and in previous experiments, only about 20% of the transcriptionally upregulated genes in whole J2 transcriptome analysis that contain signal peptide sequence and no transmembrane domain could also be confirmed to be gland localized by in situ hybridization assays [13]. As a result, whole gland transcriptomes may offer a more gland-targeted approach to effector discovery. Whole gland transcriptomes have been optimized from isolated glands for pinewood, cyst, root-knot and root-lesion nematodes, and have been useful in identifying nematode effectoromes [14–19].

Here we have undertaken a gland transcriptome analysis of *M. chitwoodi* pre-parasitic J2s. This data has allowed us to improve *M. chitwoodi* genome annotation and identify effector candidates that would be gland localized and relevant for early stages of parasitism. The gland transcriptome has shown that pre-parasitic J2s express known and putative novel effectors. We selected a few effectors (both known and novel) and confirmed their gland localization by ISH, indicating that the gland library transcriptomes can provide sets of highly confident effector candidates. By identifying and characterizing novel effectors through gland transcriptome analysis, we better understand nematode parasitism, which will ultimately facilitate developing effective management strategies for this economically damaging nematode species.

## Results

### Evidence-based genome annotation improved the existing annotation of the *M. chitwoodi* genome

To better predict *M. chitwoodi* protein-coding genes from transcriptome analyses, the first step was to improve the *M. chitwoodi* genome annotation (S1 Fig). To reannotate the genome, the *M. chitwoodi* J2 RNA-seq data [11] was combined with the newly generated *M. chitwoodi* pre-parasitic J2 gland-specific RNA-seq data (this paper). In addition, to reannotate gene structure, the sequences of proteins from Invertebrate OrthoDB [20] were used with the sequences of proteins identified from: 1) three RKN species (*M. javanica, M. arenaria, M. incognita*), 2) more distantly related PPN, *Heterodera schachtii* (cyst nematode), 3) animal-parasitic nematodes *Trichinella spiralis, Loa loa, Brugia mallayi, Necator americanus*, and 4) free-living nematodes *Caenorhabditis remanei, C. elegans, and C. briggsae*. Using this approach, a total of 12,205 *M. chitwoodi* genes were predicted. This number is similar to the 12,295 coding genes annotated for the publicly available *M. chitwoodi* genome assembly (Table 1). However, a high annotation BUSCO score of *M. chitwoodi* reannotated genome was achieved, meaning a better completeness of gene content compared to that of the publicly available genome (Table 1).

**Table 1.**
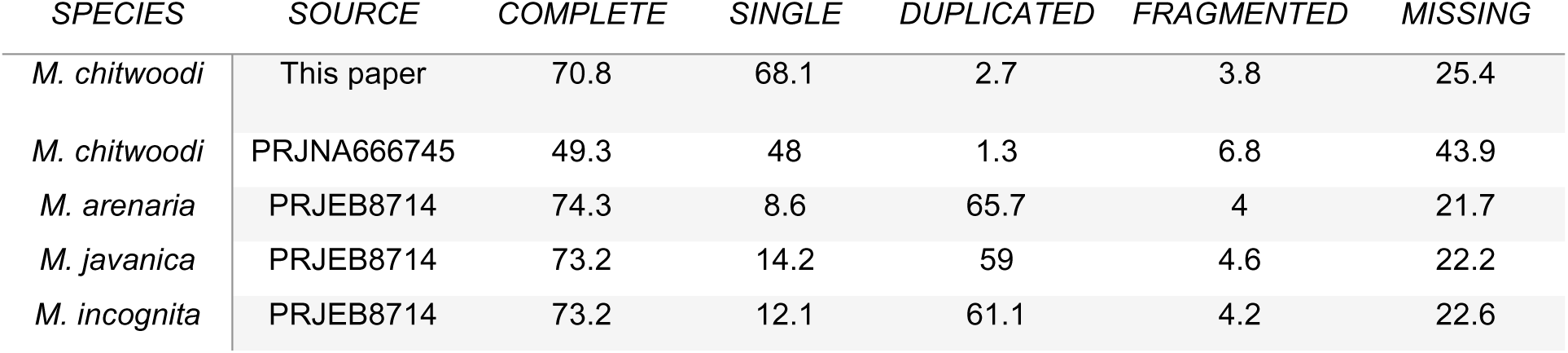
Genome annotation features for root-knot nematodes (RKNs). The table shows a comparison between previous gene annotations for RKN and the current Meloidogyne chitwoodi annotation. BUSCO scores were calculated based on eukaryota_odb10 dataset.

| SPECIES | SOURCE | COMPLETE | SINGLE | DUPLICATED | FRAGMENTED | MISSING |
| --- | --- | --- | --- | --- | --- | --- |
| *M. chitwoodi* | This paper | 70.8 | 68.1 | 2.7 | 3.8 | 25.4 |
| *M. chitwoodi* | PRJNA666745 | 49.3 | 48 | 1.3 | 6.8 | 43.9 |
| *M. arenaria* | PRJEB8714 | 74.3 | 8.6 | 65.7 | 4 | 21.7 |
| *M. javanica* | PRJEB8714 | 73.2 | 14.2 | 59 | 4.6 | 22.2 |
| *M. incognita* | PRJEB8714 | 73.2 | 12.1 | 61.1 | 4.2 | 22.6 |

To further assess the quality of our new *M. chitwoodi* annotation, a species shared and specific gene family (orthogroup) analysis was performed using Orthofinder [21]. We compared our newly reannotated *M. chitwoodi* genome to the publicly available *M. chitwoodi* and *M. arenaria* genome annotations. *M. arenaria* was chosen because it has the best publicly available RKN genome annotation, as shown by its BUSCO score (Table 1). There were 1,823 orthogroups shared between the two versions of *M. chitwoodi* annotations. However, the analysis indicated that there were 2,153 shared orthogroups between our *M. chitwoodi* genome annotation and *M. arenaria* genome annotation that were missing in the publicly available *M. chitwoodi* genome annotation (Fig 1). This indicates that our new annotation has a more complete gene family profile compared to the current available *M. chitwoodi* genome annotation.

**Fig 1.**
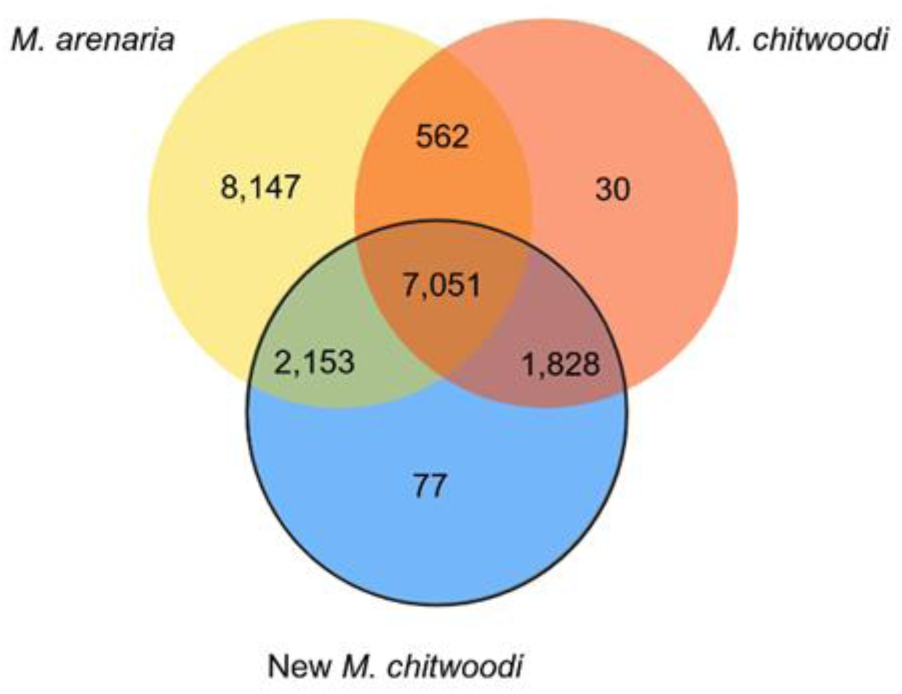
The newly re-annotated *M. chitwoodi* genome shares more orthogroups with *M. arenaria* than with the previous *M. chitwoodi* genome annotation. Venn diagram showing the shared gene families (orthogroups) between the newly re-annotated *M. chitwoodi,* current version *M. chitwoodi*, and *M. arenaria* genomes annotations.

We noticed that several putative effectors were not properly predicted by the current annotation, such as *M. chitwoodi* homologue of MiPG1, a polygalacturonase with a signal peptide and fibronectin and glycoside hydrolase 28 domains [22]. A BLASTp search for homologues using the Wormbase ParaSite resulted in the gene Mc1_00280, which is predicted to have a kinase domain (Fig 2A). Alignment of Mc1_00280 and homologues from other RKNs show that only *M. chitwoodi*’s genome predicts a kinase domain (Fig 2A). Further alignment of the gland transcriptome reads with the publicly available genome annotation shows that there are no reads present for the region comprising the predicted kinase domain (Fig 2B). The new genome annotation predicts that the region has two distinct transcripts: a polygalacturonase with its expected canonical domains (Mc1g003100.t1) and a casein kinase (Mc1g003100.t2). This shows that the polygalacturonase expressed in the gland transcriptome indeed does not include a kinase domain and that the second gene that was further annotated, the casein kinase (Mc1g003100.t2), is not expressed in the glands of pre-parasitic J2s and, for that reason, we cannot observe any reads in our gland transcriptome.

**Fig 2.**
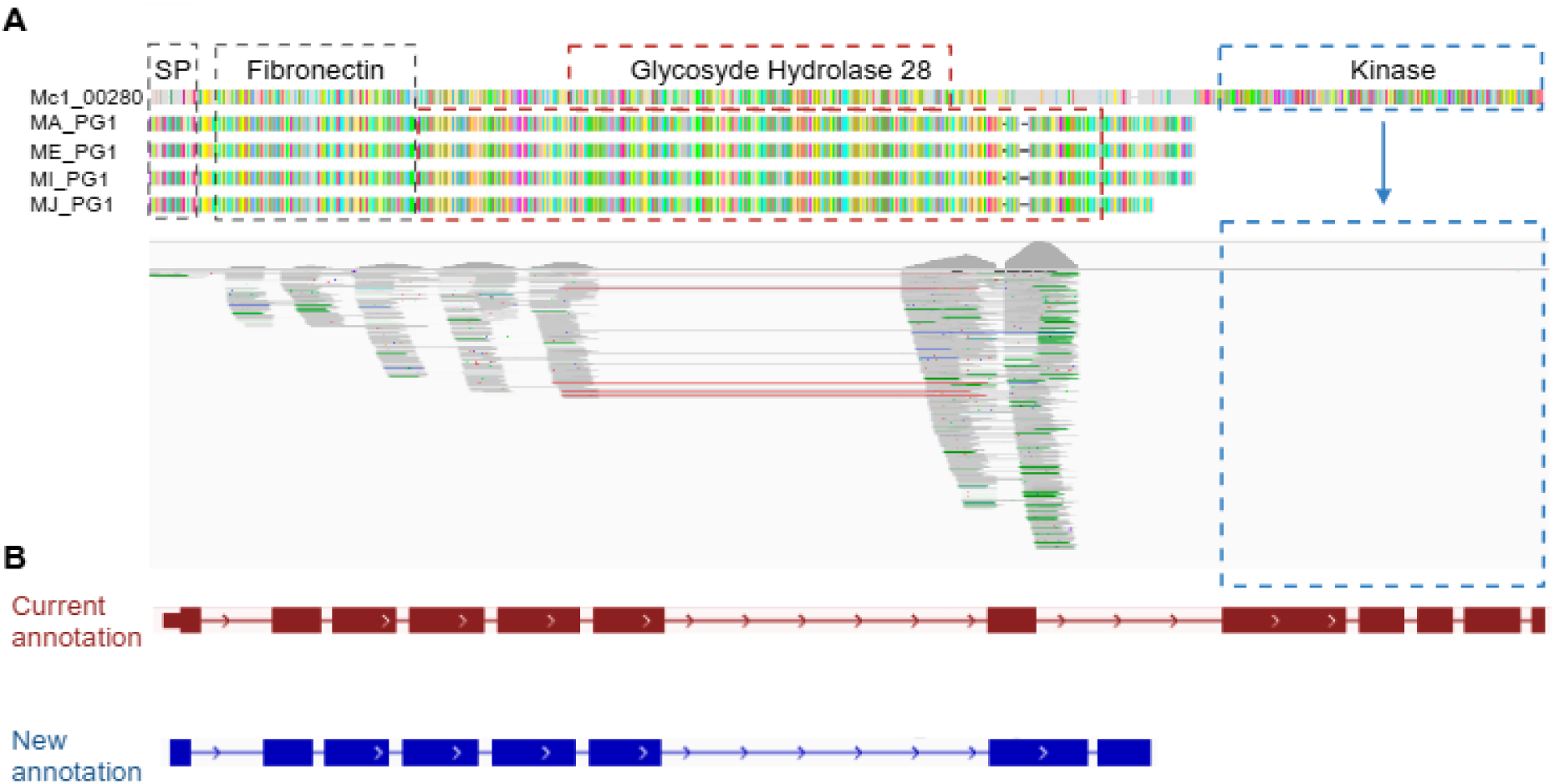
Genome reannotation improved the *M. chitwoodi* gene structure prediction. **A.** Protein alignments of polygalacturonases from *M. chitwoodi* (Mc1_00280), *M. arenaria* (MA_PG1), *M. enterolobii* (ME_PG1), *M. incognita* (MI_PG1) and *M. javanica* (MJ_PG1), showing that only *M. chitwoodi*’s current genome annotation predicts a protein with a kinase domain. **B.** Alignment of gland transcriptome reads, current genome annotation (in blue) and new annotation (in orange), validating the gene structure predicted by the new annotation. Domains are highlighted in dashed boxes and domains that differ from the others in *Meloidogyne* spp. are in red and blue. Protein alignment was performed using Genious software and transcriptome alignment to the genome was performed using IGV. Created in BioRender. Teixeira, M. (2025) https://BioRender.com/o29t552

### *M. chitwoodi* gland transcriptome analysis revealed putative effectors

Pre-parasitic J2s esophageal gland transcriptomes generated between 18.3 to 43.3 million high quality raw reads in each of three bio-replicated cDNA libraries. Around 0.7 to 6.8 million reads were mapped to the *M. chitwoodi* genome (S1 Table). The reads mapped to 5,213 transcripts present in all three biological replicates. Although the number of mapped reads appears relatively low, it is along the same percentage of mapped reads found in the *Pratylenchus* gland library [14] and may reflect: a large proportion of read mapping ribosomal genes and/or the stringency of the gland transcript analysis pipeline in mapping to the genome. Most of the plant-parasitic nematode (PPN) effectors are secreted through the canonical ER-Golgi secretory pathway and therefore, characterized by the presence of a predicted signal peptide and lack of a predicted transmembrane domain [12]. Using these criteria, 348 putative effectors were identified and represented in this first gland library generated for *M. chitwoodi* (S2 Table)

Using our new *M. chitwoodi* genome annotation, we revisited the previously published J2 transcriptome data [11]. The new analysis resulted in 10,001 transcripts identified from the whole pre-parasitic J2 transcriptome, compared to the 5,213 transcripts identified from the gland cells transcriptome. Using the criteria that effectors must have signal peptide sequence and no transmembrane domain(s), 783 possible effectors were retrieved from the J2 transcriptome (Fig 3). The comparison between the gland and the whole J2 transcriptomes revealed 337 genes in common between the gland the J2 transcriptome libraries (Fig 3, S3 Table). The 337 genes were subjected to a BLAST homology search and there were 18 genes identified that were homologous to previously described RKN effectors (S4 Table). The normalized read counts (transcripts per million, TPM) of all 18 known effectors were assessed, and Mc14g000910, a putative homolog of the known effector MiXyl1 [23], had the lowest read count from the gland transcriptome. The expression of this gene was used as the threshold of minimum effector gene expression. From the initial 337 putative effectors, 111 effector candidates (S5 Table) passed the following criteria: 1) gene expression is above the minimum expression level set by Mc14g000910; 2) encode proteins with predicted secretion signal sequences; 3) encode proteins with no predicted TM domains. Among these 111 candidate effectors, 72 contained known protein domain sequences, including sequences for calreticulin, glycoside hydrolase, c-type lectin-like, peptidase, which are protein domains found in plant-parasitic nematode effectors (S2 Fig). Thirty-nine out of 111 candidate effectors have no identified domain(s) based on Interpro or CDD (NCBI Conserved Domain Database) searches (S6 Table). Twenty-six had protein sequences with more than 50% identity to proteins from *M. graminicola* or *M. enterolobii*, or both. The remaining 13 had no significant similarity to any proteins from other organisms, indicating they may be *M. chitwoodi* specific (S6 Table). The 13 *M. chitwoodi* specific putative effectors were further analysed by comparing number of reads for each in the pre-parasitic J2s and the parasitic J2 [11]. From these 13 putative effectors, 7 were more highly expressed in the pre-parasitic J2 stage compared to the parasitic life stage (8 dpi in potato roots) (Table 2)

**Fig 3.**
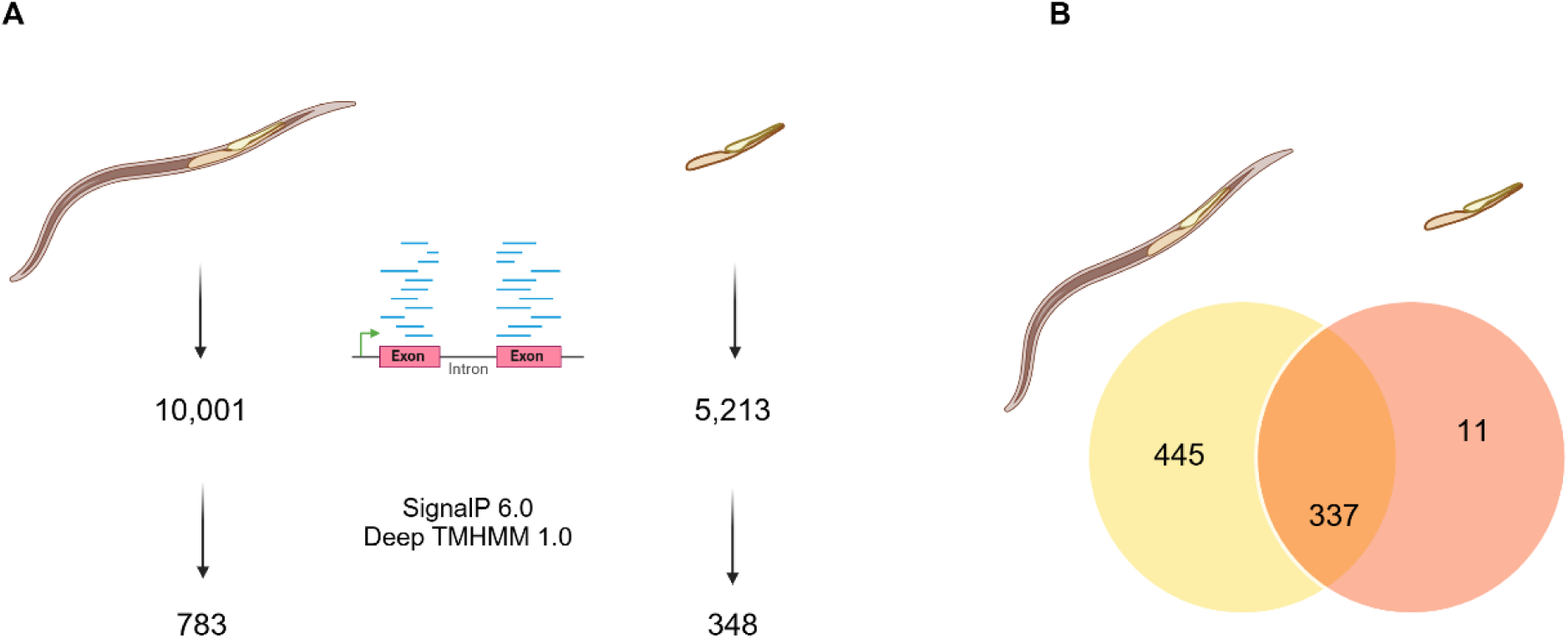
Transcriptome analysis summary. **A.** Comparison between the number of transcripts retrieved from whole J2s and their glands. **B.** Venn diagram showing number of overlapping putative effectors identified in pre-parasitic J2 and their glands. Created in BioRender. Teixeira, M. (2025) https://BioRender.com/f42i870

**Table 2.**
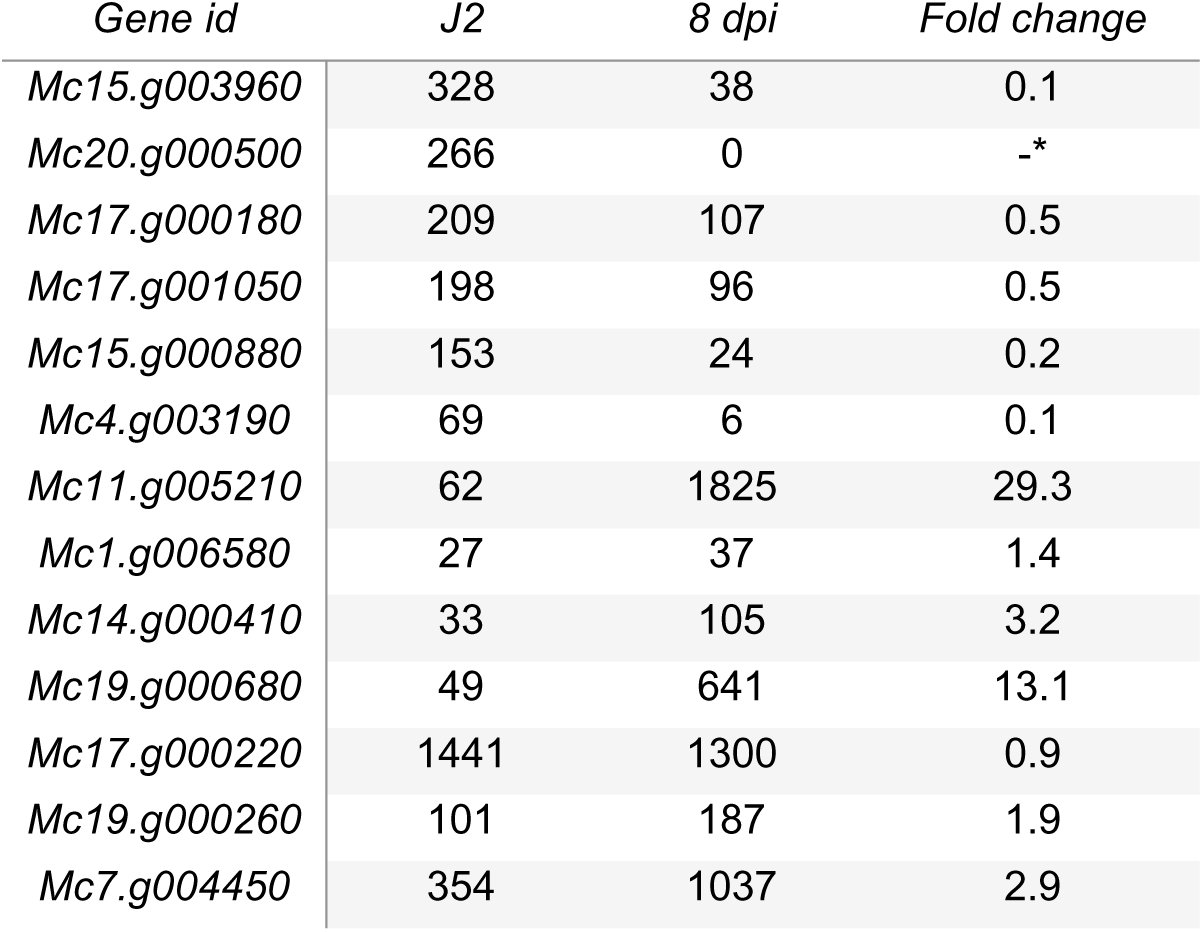
Transcripts of putative *Meloidogyne chitwoodi* specific effectors, with unknown domains. Table shows transcripts per million (TPM) from pre-parasitic J2s and for parasitic nematodes at 8 dpi. Fold change was calculated relative to pre-parasitic J2 TPMs. *Transcript Mc20.g000500 was not expressed at 8dpi.

To confirm our *in-silico* analysis of the gland library, the expression of 6 genes was quantified by qRT-PCR in three life stages (eggs, pre-parasitic J2s and parasitic nematodes at 4 dpi) (Table 2). First, we looked at gene expression of three known effectors identified in our gland library: Mc5g006680 and Mc1g003100, which are glycoside hydrolases with high similarity to MiENG1 [24] and MiPG1 [22], respectively, and Mc5g002180, which has similarity to the described venom allergen protein Mi-msp- 1 [25]. Next, we looked at the expression of Mc11229, a proteinase-inhibitor like, which was previously characterized as a gland localized effector from *M. chitwoodi* [26]. Lastly, we measured gene expression for two putative effectors from the gland library analysis. One was an effector called Mc4g000590 that had putative homologs in *M. graminicola* and *M. enterolobii*. The second was Mc15g003960, which was *M. chitwoodi* specific based on BLAST analysis and shows one of the highest fold change differences in the comparison between the pre-parasitic J2s and the parasitic stage (8 dpi in potato roots) (Table 2).

All 6 genes whose expression was investigated by qRT-PCR were highly expressed in pre-parasitic J2 stage compared to the egg stage (Fig 4). The three known effector genes (the two glycoside hydrolases [22, 24] and a Mi-msp-1 [25] exhibited the highest level of expression in parasitic nematodes (4 dpi in potato) compared to eggs and pre-parasitic J2s. Transcript abundance of two novel effectors, Mc4g000590 and Mc15g003960, and the previously identified effector Mc11229 were significantly higher in pre-parasitic J2s relative to the eggs and parasitic nematodes (4 dpi).

**Fig 4.**
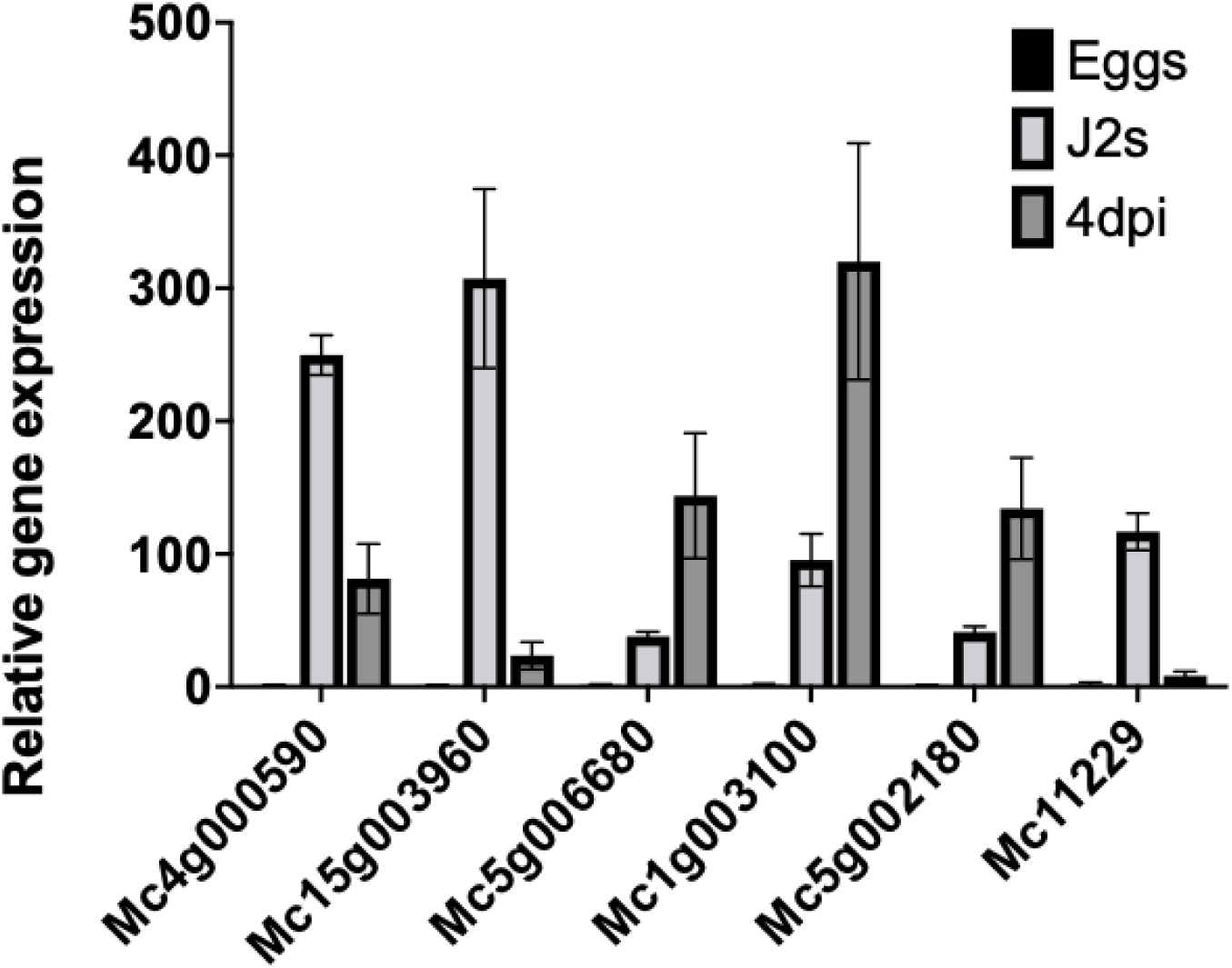
Relative expression of selected effector candidate genes at three nematode life stages (egg, pre-parasitic J2 and parasitic J2 at 4 dpi). Data is presented as the mean of fold change of two biological replicates +/- standard deviation (SD).

To further validate the gland library, ISH were performed on these six genes (Table 3, Fig 5). The anti-sense probes for all six genes localized to the esophageal gland region of the J2, making them “candidate effectors”. To test the stringency of our effector discovery method, we tested two potential effectors (presence of SP, no TM domain) that did not pass our threshold of expression in gland tissue, Mc6g004130 and Mc8g001490. The ISH showed these transcripts were not localized to the glands, i.e., Mc6g004130 localized to the hypodermis and Mc8g001490 localized to two punctate structures posteriorly to the nerve ring (Fig 5). Thus, only the putative effectors that pass our gland specific criteria were proven to be expressed in the glands, validating the gland transcriptome data and our effector discovery method.

**Fig 5.**
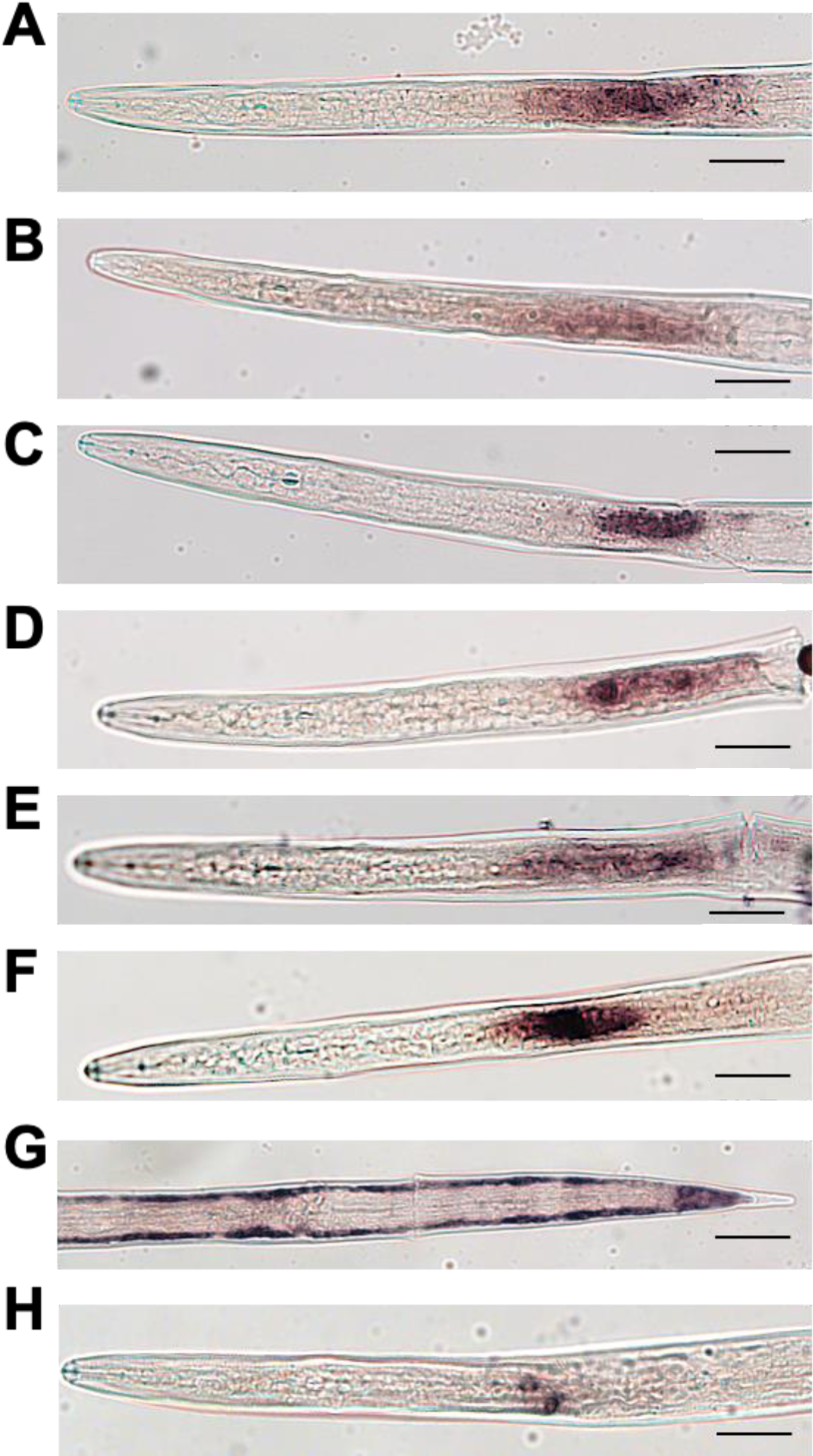
Representative photos of *in situ* hybridizations of a digoxigenin-labeled antisense probes for transcripts expressed in the J2s A-F, gland localized effectors. **A**. scaffold4.g000590 (novel *M. chitwoodi* effector, with putative homologs in *M. enterolobii* and *M. graminicola*), **B.** scaffold15.g003960 (*M. chitwoodi* specific), **C.** scaffold5.g006680 (putative glycosyl hydrolase with high similarity to MiENG1), **D.** scaffold1.g003100 (putative glycosyl hydrolase with high similarity to MiPG1), **E.** scaffold17.g005060 (effector with a cysteine-rich secretory proteins, antigen 5, and pathogenesis-related 1 proteins domain, with similarity to MiVAP), **F.** scaffold1.g000780 (described by Roze et al. 2018). **G.** scaffold6.g004130 and **H.** scaffold8.g001490, transcripts that didn’t pass our effector criteria for gland expression.

**Table 3.**
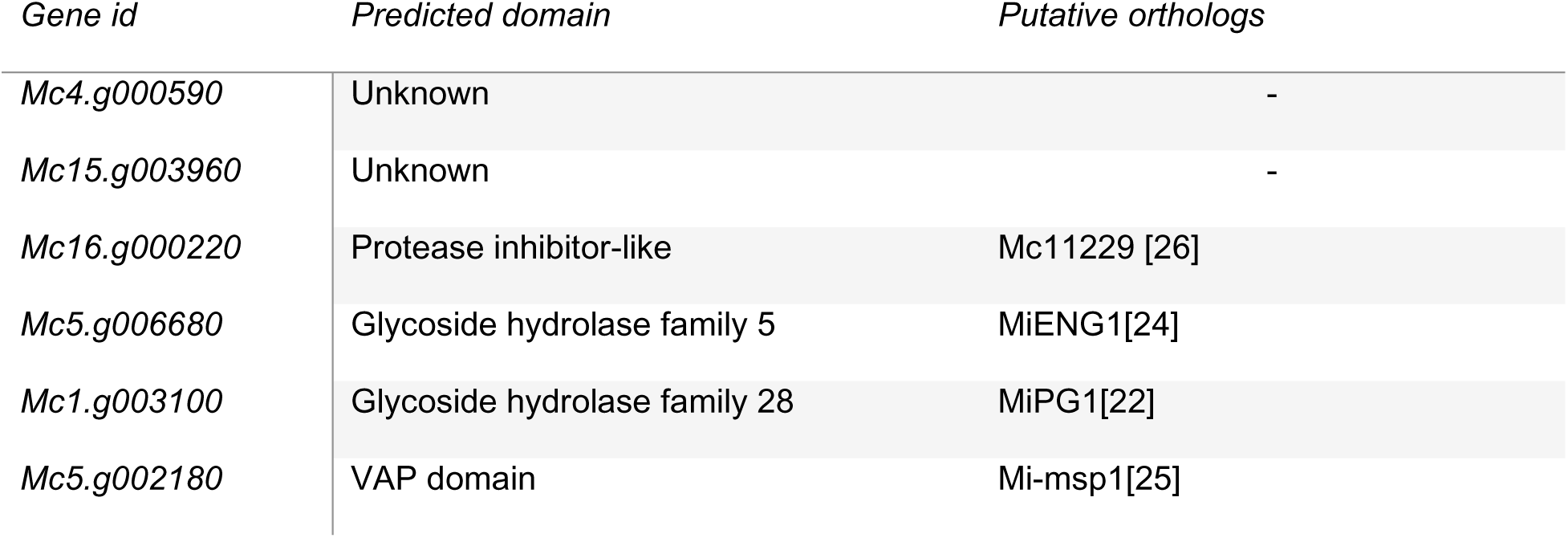
Transcripts chosen for further characterization. Domains were predicted using Interpro and NCBI Conseved Domain Database. Putative orthologs were obtained performing BLASTn searches on NCBI and WormBase parasite.

| Gene id | Predicted domain | Putative orthologs |
| --- | --- | --- |
| Mc4.g000590 | Unknown | - |
| Mc15.g003960 | Unknown | - |
| Mc16.g000220 | Protease inhibitor-like | Mc11229 [26] |
| Mc5.g006680 | Glycoside hydrolase family 5 | MiENG1[24] |
| Mc1.g003100 | Glycoside hydrolase family 28 | MiPG1[22] |
| Mc5.g002180 | VAP domain | Mi-msp1[25] |

### The novel effector, Mc15g003960, facilitated *M. chitwoodi* parasitism

Mc15g003960 was further investigated because it was discovered as a *M. chitwoodi* specific effector in the gland transcriptome analysis and confirmed to be gland localized by ISH. Moreover, the gene is more highly expressed in the pre-parasitic J2 stage compared to other life stages, as found in our whole J2 transcriptome (Table 2) and in the qRT-PCR expression analyses (Fig 4).

*Mc15g003960* is a 438 bp coding sequence that encodes a 145 amino acid protein. The signal peptide sequence cleavage site was predicted to be between the amino acids 26 and 27. To characterize its role in *M. chitwoodi* parasitism, two independent Arabidopsis transgenic lines were generated that expressed *Mc15g003960* without its signal peptide *(Mc15g003960Δsp),* driven by the CaMV35S promoter. The transformation events in the two transgenic lines were first confirmed with genomic PCR and the expression of the *Mc15g003960* in the two lines was measured by qRT-PCR (S1 Fig). The two independent Arabidopsis lines expressing *Mc15g003960Δsp* were infected with *M. chitwoodi* and assessed at 15 dpi for root galling. The fresh shoot and root weight of the two transgenic lines showed no significant difference as compared to wildtype plants (S3 Fig), suggesting that the expression of *Mc15g003960Δsp* has no effects on plant growth or development. Remarkably, both transgenic lines exhibited more galling compared to the wildtype plants at 15 dpi (Fig 6).

**Fig 6.**
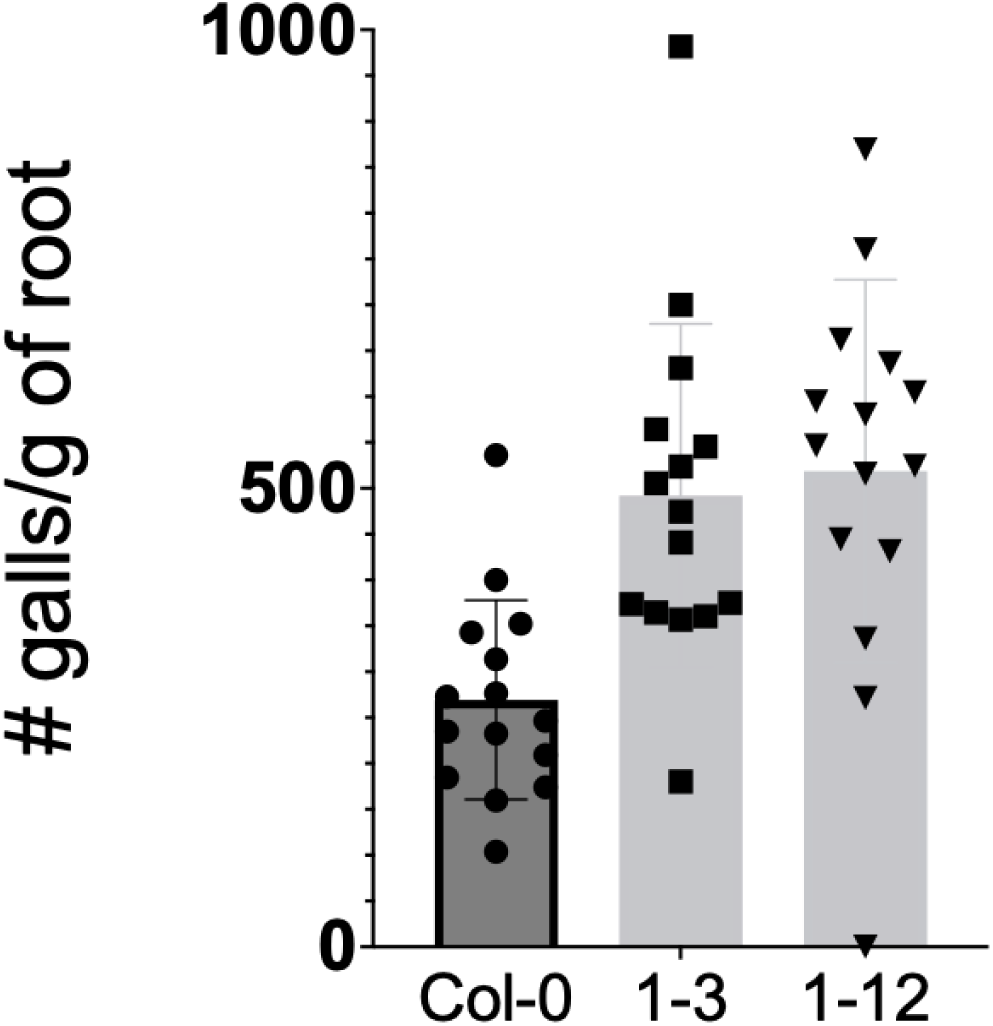
Expression of Mc15g003960 in Arabidopsis makes the plant more susceptible to *M. chitwoodi* infection. Two independent lines of Arabidopsis expressing Mc15g003960 (1.3 and 1.12) and the wildtype control plants (Col-0) were inoculated with *M. chitwoodi* Race 1 J2s. At 14 dpi, galls per plant were counted at 14 dpi. Data represents the mean number of galls per plant +/- SD (Mann-Whitney test, error bars represent SD, ns: not significant p-value>0.05, ***p-value ≤0.001, ****p-value ≤0.0001). N= 15. Experiment was repeated three times with similar results.

We hypothesized that the transgenic lines expressing *Mc15g003960Δsp* showed enhanced susceptibility to *M. chitwoodi* infection because the effector might be suppressing PTI.

We had previously identified a RKN effector that can block defense related callose deposition [27]. Callose deposits in leaves can be quickly triggered by the bacterial defense elicitor flg22 [28]. To determine if *Mc15g003960* would affect flg22-induced callose deposition in leaves, flg22 was infiltrated into the leaves of wildtype and *Mc15g003960Δsp* transgenic lines. Subsequently, flg22-induced callose deposits were detected using aniline blue staining. The leaves from both *Mc15g003960Δsp* transgenic lines showed similar levels of callose deposits when compared with the wildtype after flg22 treatment. These results suggest that flg22- triggered callose response in *Arabidopsis* leaves was not affected by the constitutive expression of *Mc15g003960* (Fig 7).

**Fig 7.**
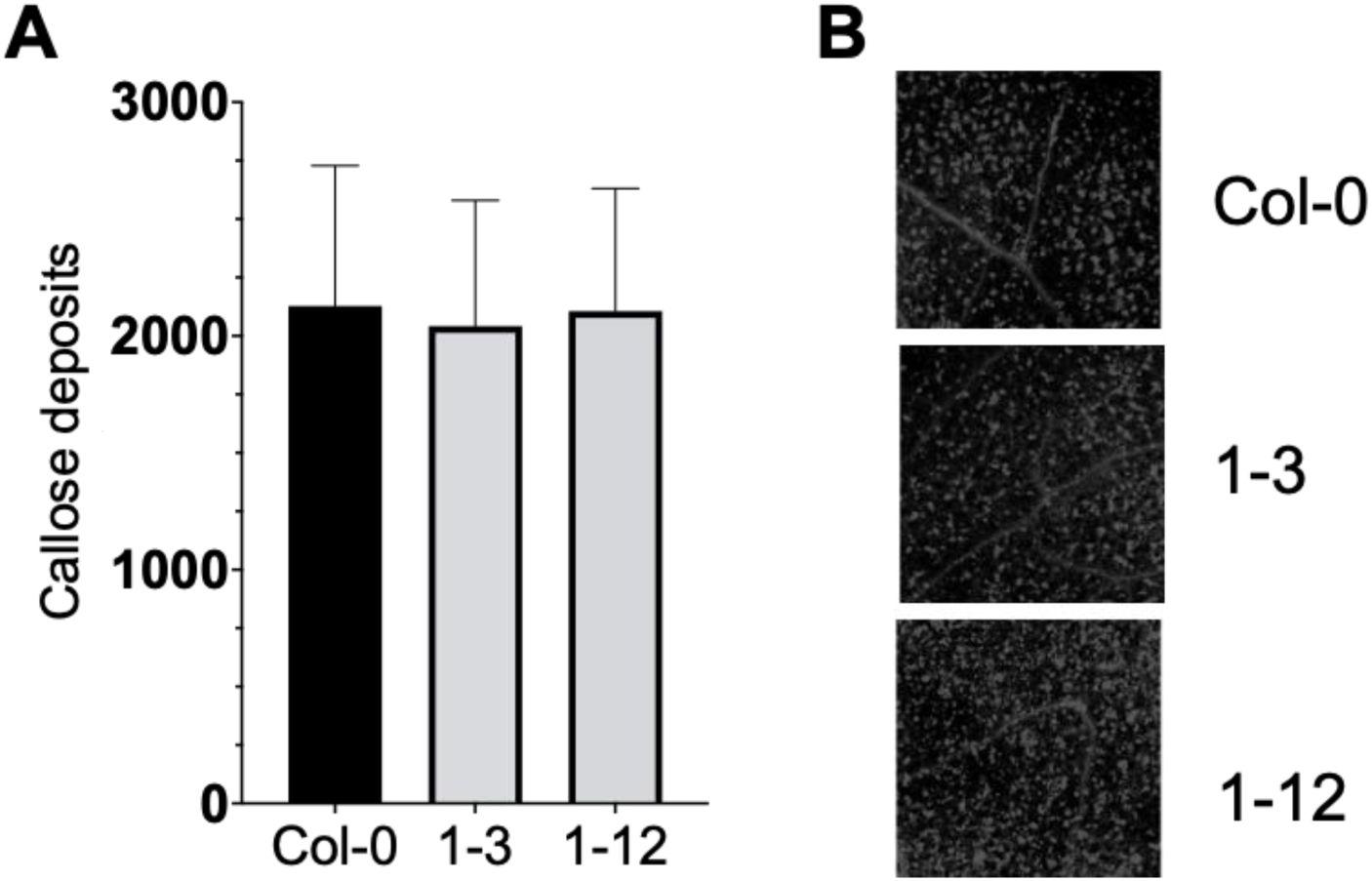
Mc15g003960 does not suppress callose deposition triggered by flg22. **A.** Quantification of aniline blue-stained callose deposits in the leaf tissue of the *Arabidopsis* plants expressing Mc15g003960 and wild-type plants after flg22 infiltration, calculated using ImageJ software. **B.** Aniline blue staining shows that callose deposition was equivalent in the leaves of transgenic *Arabidopsis* plants expressing Mc15g003960 compared with wild-type plants after flg22 treatment. (n>25).

## Discussion

Proper prediction of protein-coding genes in nematode genomes is essential for effector identification and subsequent characterization. Thus, generating new transcriptome data provides ample opportunity for enhancing the genome annotation of plant-parasitic nematodes. Among RKNs, *M. arenaria* (ASM90000398v1) currently has the best annotated genome for a published genome based on a BUSCO score of 74.3%, while *M. chitwoodi* (ASM1518303v1) presents the lowest BUSCO score of 49.3% (Table 1). Improving the genome annotation results in adequate gene structure prediction. A search for effectors using the current annotation indicated incorrect prediction for a few well characterized effectors from other RKNs. The genome re-annotation can benefit from both transcriptome and proteome data, which confer more robustness to annotation prediction [29]. Therefore, to improve *M. chitwoodi* genome annotation, we combined our previously published J2 transcriptome and the current gland transcriptome datasets, with protein datasets from other PPNs, animal-parasitic nematodes, and free-living nematodes. Although the number of predicted genes, 12,205, remains very similar to the current annotation (12,295), we achieved a better prediction of gene structure, as shown by the new BUSCO score of 70.8%. The quality of this updated *M. chitwoodi* annotation was further assessed by comparing the othogroups of two annotation versions with those of the high-quality *M. arenaria* genome. The newly annotated genome identified 2,153 orthogroups between *M. chitwoodi* and *M. arenaria* that were absent in the previous version, indicating a more complete *M. chitwoodi* genetic makeup.

The whole pre-parasitic J2 transcriptome analysis indicated that there were 783 possible effectors, following the criteria of coding for proteins with a signal peptide sequence and no transmembrane domains. However, it is important to note that the majority of known effectors are mainly produced in the nematode esophageal glands and secreted via the stylet [12]. Therefore, focusing on genes expressed within the nematode glands would be a more efficient way to identify putative effectors relevant for successful parasitism. For instance, gland transcriptomes generated for other nematode species, such as *Globodera rostochiensis*, *Pratylenchus penetrans, Radopholus similis, Bursaphelenchus xylophilus*; *M. incognita*, *H. schachtii*, and *H. glycines* [14, 15, 17, 18, 30–32], reveal large sets of highly confident effector candidates. In the case of *M. chitwoodi*, 337 of the 783 transcripts coding for secreted proteins were found in both the whole J2 and the gland transcriptomes. This suggests that, akin to other PPNs, the transcriptome of the glands library offers a more refined and targeted pool of potential effector candidates. By focusing on this specific subset, it allows for a more precise identification of key molecular players, providing a higher level of specificity compared to broader approaches. It should be noted that not being gland specific does not preclude genes from being effectors as they may be secreted from nematode organs other than the stylet.

A more conservative criterion was applied by considering only genes with expression levels exceeding that of the known effector with the lowest expression level, leading to a final set of 11 putative genes. These represent around 2% of the 5,213 total transcripts initially retrieved from our gland transcriptome. Other publications found that putative effectors represent 2.42% [30] and 7.5% [31] in *M. incognita* glands, and 2% in *P. penetrans* gland transcriptomes [14]. These variations might reflect the different bioinformatic strategies deployed for effector discovery. Our approach used known effectors in RKN to define a minimum level of effector gene expression. This stringent filtering step may exclude true effectors that have expression lower than this cutoff but allowed us to narrow our list of effectors to a more manageable number of candidates.

Among our final candidates, the majority (65%) encode proteins with known domains, many of which have been found in other RKN species. The remaining transcripts (35%) are novel proteins with unknown domains, from which 12% encode *M. chitwoodi* specific proteins and 23%, proteins with similarity to *M. enterolobii* or *M. graminicola*, or both. Thus, our results show a small proportion of candidate genes encoding pioneer sequences, which suggests that most of the effectors were present before diverging from the last common ancestor, unlike *Pratylenchus*, which contains a large number (57%) of unique effectors [14].

To characterize novel effectors from *M. chitwoodi* involved in the early stages of parasitism, we analyzed their expression using our previously published transcriptome data from whole pre-parasitic and parasitic J2s [11]. Among the thirteen *M. chitwoodi*- specific effectors, six were found to be highly expressed in pre-parasitic J2s compared to parasitic nematodes. Of these, Mc15g003960 was selected for further study. When expressed in *Arabidopsis*, Mc15g003960 increased the plant’s susceptibility to *M. chitwoodi*. While many root-knot nematode effectors are known to suppress plant defense responses, Mc15g003960 did not inhibit flg22-induced callose deposition, suggesting it may target other aspects of PTI or perform additional roles in facilitating nematode-plant interactions. Future studies will further explore Mc15g003960’s role in nematode parasitism and identify its interacting plant proteins.

## Materials and Methods

### Reannotation of *M. chitwoodi* genome

*M. chitwoodi* softmasked genome (ASM1518303v1) was obtained from Wormbase ParaSite database under Bioproject ID: PRJNA666745 [5]. Genome quality was assessed using eukaryote_odb10 dataset [32]. The genome annotation was conducted at the high-performance computer cluster “Kamiak”, at Washington State University and a pre-built genome structural and functional annotation workflow [33] was adapted in this study. In brief, GEMmaker (v2.1.0) [34] was used to map *M. chitwoodi* gland transcriptome generated from this study and previously generated *M. chitwoodi* J2 RNA-seq data [11] to the softmasked genome. The generated bam files were used as hint files for BRAKER1 [35] an RNA-seq based genome structural annotator. As suggested by Gabriel *et al.* 2021 [36], the annotation accuracy can be enhanced by running the protein homology-based genome annotator BRAKER2 [37]. Therefore, Invertebrate OrthoDB [20], *Heterodera schachtii, M. arenaria, M. javanica, M. incognita, Trichinella spiralis, Loa loa, Brugia mallayi, Necator americanus*, *Caenorhabditis remanei, C. elegans, and C. briggsae* protein datasets were used as hints for BRAKER2 to annotate *M. chitwoodi* genome. Later, TSEBRA [36] was used to combine the BRAKER1 and BRAKER2 results. To further improve the annotation, EviAnn was used to annotate genes purely based on the same set of protein and rna-seq data used previously. Bedtool2 intersect [38] was used to combine the BRAKER, EviAnn, and previous annotation [5]. The final structurally annotated *M. chitwoodi* genome assembly was fed into EnTAPnf [39] for gene functional annotation with gene ontology assignment. The completeness of functionally annotated genome assembly was assessed by BUSCO using eukyrote_odb10 dataset [32]. The enrichment of gene expression in the gland transcriptome was visualized using Integrative Genomics Viewer (IGV) software (v. 2.10.3) [40]. The species-specific gene family (orthogroup) analysis was adapted from Zhang et al. (2022) [21]. Orthofinder [41] was used to assign each gene to a protein family.

### Transcriptomic analysis

Five to 10 esophageal glands were extracted from pre-parasitic second stage juveniles of *M. chitwoodi* race 1 and pooled to form one sample [42]. Libraries were prepared using Takara Bio’s SMART-seq mRNA LP (634768) kit, which includes RNA isolation and library preparation. Three libraries were sequenced by Novogene, using Illumina Novaseq 6000 (PE 150bp with 250M read depth).

Raw sequencing data were process using the gene expression analysis workflow GEMmaker (v2.1.0) [34]. STAR aligner implemented in GEMmaker was used to map the transcriptomes to the *M. chitwoodi* race 1 draft genome [5]. The resulting sequence mapping quality report and normalized gene count matrix were used in this study.

### Prediction of putative effectors

For possible effector gene identification from all transcripts, signal Peptides were predicted using SignalP6.0 [43, 44] and transmembrane domains were predicted using DeepTMHMM1.0 [45], both using default settings. Domain prediction was performed using Interpro [46] and NCBI’s conserved domain database [47], using amino acid sequences. To identify homolog of candidate effector gene, BLASTp and BLASTn search was carried out at the NCBI online servers using non-redundant protein dataset and the expressed sequence tags (ESTs) database respectively using the default parameter. Manual curation of putative effector sequences was performed aligning the gland transcriptome data to the genome annotation using Integrative Genomics Viewer (IGV) software v. 2.10.3) [42].

### Cloning and transgenic lines generation

Mc15g003960 coding sequence, without its predicted signal peptide sequence (*Mc15g003960Δsp*), was amplified from *M. chitwoodi* race 1 cDNA by PCR (S7 table) and inserted into the Gateway donor vector pDONR207. It further was recombined into Gateway destination vector pB2GW7 to produce 35S::*Mc15g003960Δsp*.After sequencing, the plasmid was used to transform *Agrobacterium tumefaciens* GV3101, which was used to transform Arabidopsis Col-0 following the floral dip protocol of Clough and Bent (1998) [48]. Homozygous T3 seeds from two independent transgenic lines were collected from T2 lines after segregation analysis on BASTA-containing medium. Nematode gene expression in the transgenic plants was confirmed by qPCR (S7 Table).

### Nematode cultures

*M. chitwoodi* race 1 (initially provided by Dr Charles Brown from USDA-ARS, Prosser, WA) was multiplied on susceptible tomato *Solanum lycopersicum* cv. Rutgers under greenhouse conditions. Egg mass collection and J2 hatching was performed as previously described [11].

### Nematode infection assays

Arabidopsis seeds were surface-sterilized and sown on Murashige & Skoog (Duchefa) agar plates (½MS salts, 1% sucrose, 0.8% agar, pH 6.4). Three week old seedlings were transferred to 500 ml sand-filled cone-tainers kept in growth chamber (14 h:10 h light:dark cycle, 23 °C). One week after transplanting to sand, the plants were inoculated with 750 *M. chitwoodi* eggs per plant. At 15 days post-inoculation, the plants were gently removed from the cone-tainers. Each root system was weighed and the number of galls per plant was assessed. Nematode infection assays were repeated twice with similar results.

### RNA extraction and Real time PCR validation

Susceptible Russet Burbank plants were propagated in tissue culture for three weeks and then transferred to 500 ml cone-tainers filled with sand and subsequently maintained in growth chambers at 14 h:10 h light:dark cycle, 23 °C. Fourteen days after transplanting to sand, the plants were inoculated with 500 freshly hatched *M. chitw*oodi J2s. Samples were collected 4 days post inoculation (dpi) and immediately frozen for RNA extraction.

For gene expression validation, two biological samples of eggs, pre parasitic J2s and galled potato root tissue at 4 dpi were collected. Total RNA was extracted using the Trizol method. Briefly, trizol was added to ground sample and RNA was precipitated overnight using isopropanol. RNA was then purified using ethanol and resuspended in DEPC water. 1ug of RNA was treated with DNA-freeTM DNA Removal Kit (ThermoFisher) to remove residual DNA and ProtoScript II First Strand cDNA Synthesis Kit (New England Biolabs, USA) was used to prepare cDNA. Quantitative PCR was performed using SsoAdvancedTM Universal SYBR Green Supermix on a CFX96 Real-Time PCR Detection System (Bio-Rad, USA) and the primers listed in S7 Table. The qRT-PCR conditions were: 95C for 3 min, 40 cycles for 95C for 15 sec, 53C for 15sec and 72C for 20 sec, followed by a melting curve analysis from 65C to 95C with 0.5C increments at 5sec. Each experiment consisted of three technical replicates per sample.

Expression levels of genes of interest in nematode were normalized to the expression of Internal transcribed spacer 2 (ITS2) rRNA, a housekeeping gene from *M. chitwoodi* [50]. The relative expression levels of each target gene from pre parasitic J2s and 4 dpi tissue were calculated by comparing with those in eggs using the 2–ΔΔCt method [49].

### Callose deposition assay

Leaves from 4-week-old Arabidopsis plants were infiltrated with 1uM flg22. After 24h, leaves were punched using a cork borer and fixed in 95% ethanol overnight. Samples were then rehydrated with a serial incubation with 50% ethanol for 30 minutes, followed by two 30 min incubations with a phosphate-buffered saline, stained with 0.8% aniline blue for 1h and visualized under an AxioObserver A1 inverted microscope using a DAPI filter. Callose deposition was quantified with ImageJ [50]. At least 25 leaf punches were collected from four plants per genotype using a 4 mm cork borer, all images were taken at 10X magnification and the experiment was repeated twice.

### In situ hybridization

Whole mount in situ hybridizations were performed for *M. chitwoodi* J2 following the protocol of de Boer et al. [51]. Specific PCR primers were used to amplify the genes of interest from *M. chitwoodi* J2 cDNA (S7 Table). The resulting PCR products were then used as a template for generation of sense and antisense DIG-labeled probes using a DIG-nucleotide labeling kit (Roche). Hybridized probes within the nematode tissues were detected using an anti-DIG antibody conjugated to alkaline phosphatase and its substrate. Nematode segments were observed using a Nikon Eclipse 5*i* light microscope.

### Statistics

Statistical tests were performed using GraphPad Prism version 8.00 for Windows (GraphPad Software, La Jolla California, USA).

## Acknowledgements

All computation works were performed using Dr. Stephen Ficklin’s “Computational Biology Lab” node resource and “Kamiak” HPC at Washington State University, US.

## Data Availability

The *M. chitwoodi* RNA-seq data have been deposited at the NCBI database under BioProject ID PRJNA1190951 and the M. chitwoodi genome re-annotation is found at **10.5281/zenodo.14542289.**

## Supporting Information

**S1 Fig. A schematic workflow of the genome annotation pipelines used in this study**. Teixeira, M. (2025) https://BioRender.com/f18v715

**S2 Fig. Domain prediction of proteins encoded by putative effectors.** Prediction performed using Interpro and NCBI Conserved Domain Database.

**S3 Fig. Evaluation of two independent lines expressing Mc15g003960 (1.3 and 1.12). A.** transgene expression from each line. **B**. Root weight of the two lines and Col-0 at 14 dpi. Data show mean weight of the roots +/-SD. (Mann Whitney test, error bars represent SD, ns: not significant). N=15. Experiment was repeated three times with similar results.

**S1 Table.** Summary of Gland transcriptome sequencing and alignment data.

**S2 Table**. Gland transcripts with SP and no TM domain

**S3 Table**. Common transcripts between glands and J2

**S4 Table**. Putative homologs of known effectors

**S5 Table.** Putative Mc1 effectors after using homologs as cut-off criteria.

**S6 Table**. Transcripts for which no domain was predicted using Interpro and NCBI Conserved Domain Database.

**S7 Table.** Primers used in this paper

