## Supplementary material for "Genome reannotation and effector candidate identification in *Meloidogyne chitwoodi* through gland-specific transcriptome analysis": Gland paper S figures_04.02.25.pdf

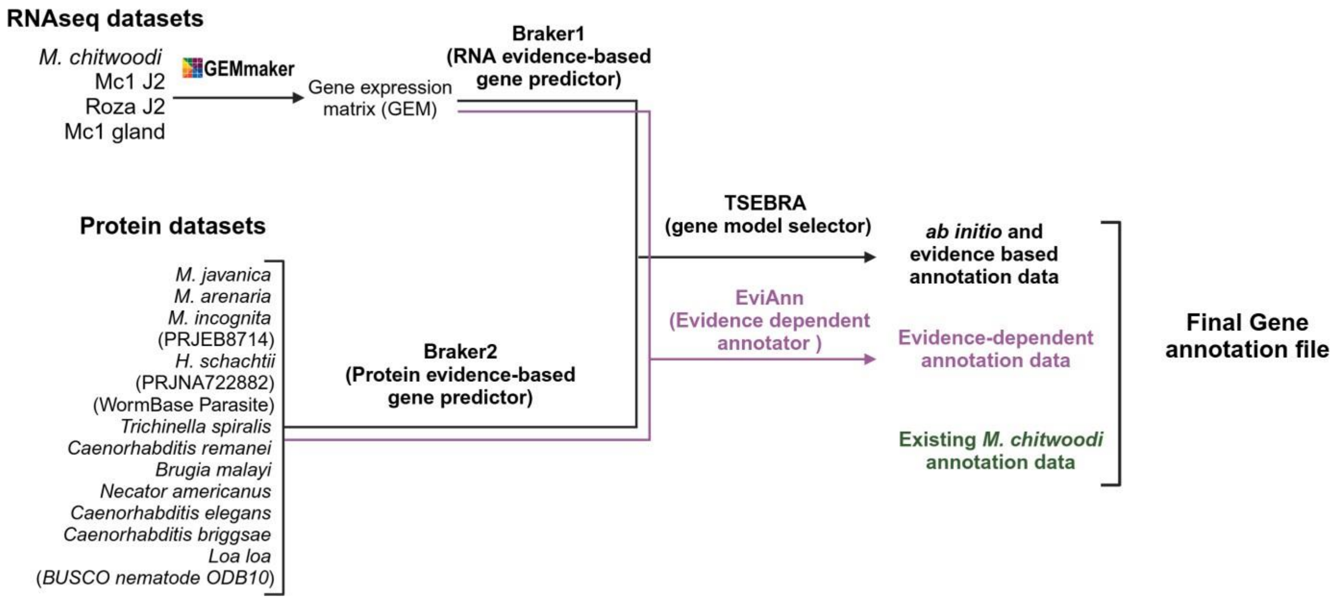


**S1 Fig. A schematic workflow of the genome annotation pipelines used in this study**. Teixeira, M. (2025) https://BioRender.com/f18v715

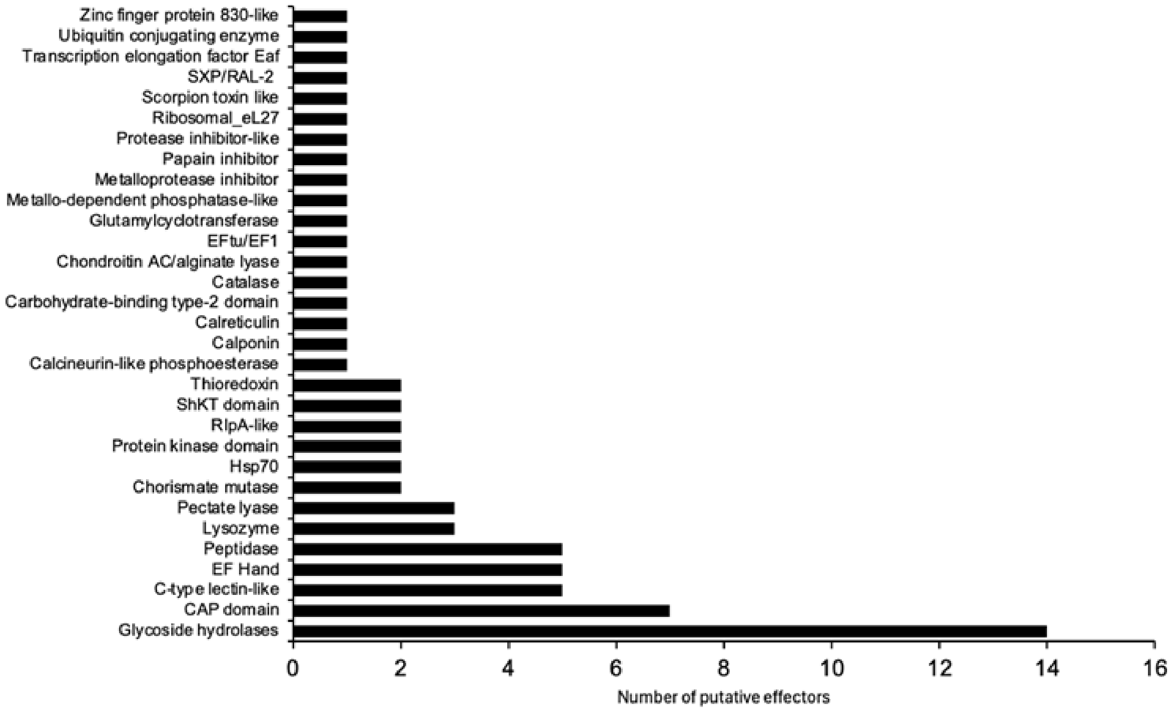


**S2 Fig. Domain prediction of proteins encoded by putative effectors.** Prediction performed using Interpro and NCBI Conserved Domain Database.

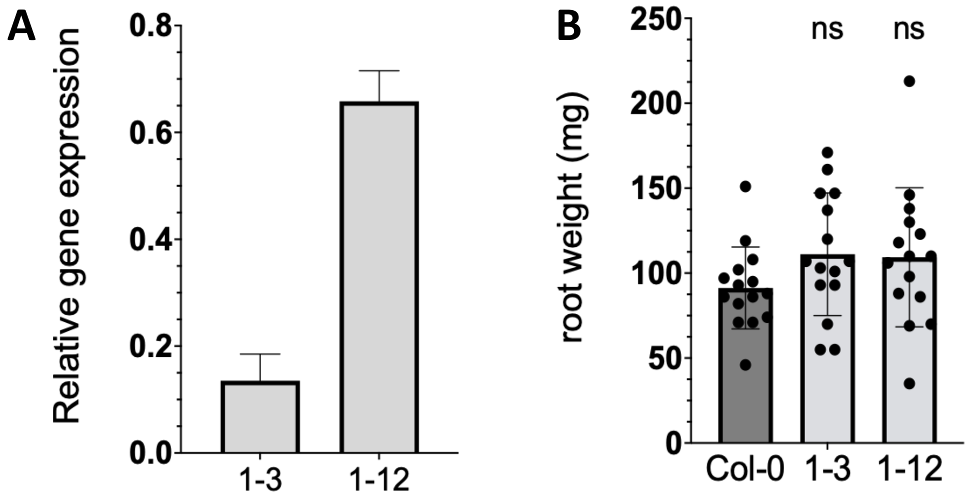


**S3 Fig. Evaluation of two independent lines expressing Mc15g003960 (1.3 and 1.12). A.** transgene expression from each line. **B**. Root weight of the two lines and Col-0 at 14 dpi. Data show mean weight of the roots +/-SD. (Mann Whitney test, error bars represent SD, ns: not significant). N=15. Experiment was repeated three times with similar results.
